# White Matter Microstructure and its Relation to Clinical Features of Obsessive-Compulsive Disorder: Findings from the ENIGMA OCD Working Group

**DOI:** 10.1101/855916

**Authors:** Fabrizio Piras, Federica Piras, Yoshinari Abe, Sri Mahavir Agarwal, Alan Anticevic, Stephanie Ameis, Paul Arnold, Núria Bargalló, Marcelo C. Batistuzzo, Francesco Benedetti, Jan-Carl Beucke, Premika S.W. Boedhoe, Irene Bollettini, Silvia Brem, Anna Calvo, Kang Ik Kevin Cho, Sara Dallaspezia, Erin Dickie, Benjamin Adam Ely, Siyan Fan, Jean-Paul Fouche, Patricia Gruner, Deniz A. Gürsel, Tobias Hauser, Yoshiyuki Hirano, Marcelo Q. Hoexter, Mariangela Iorio, Anthony James, Janardhan Reddy, Christian Kaufmann, Kathrin Koch, Peter Kochunov, Jun Soo Kwon, Luisa Lazaro, Christine Lochner, Rachel Marsh, Akiko Nakagawa, Takashi Nakamae, Janardhanan C. Narayanaswamy, Yuki Sakai, Eiji Shimizu, Daniela Simon, Helen Blair Simpson, Noam Soreni, Philipp Stämpfli, Emily R. Stern, Philip Szeszko, Jumpei Takahashi, Ganesan Venkatasubramanian, Zhen Wang, Je-Yeon Yun, ENIGMA OCD Working Group, Dan J. Stein, Neda Jahanshad, Paul M. Thompson, Odile A. van den Heuvel, Gianfranco Spalletta

## Abstract

**Importance:** Microstructural alterations in cortico-subcortical connections are thought to be present in Obsessive-Compulsive Disorder (OCD). However, prior studies have yielded inconsistent findings, perhaps because small sample sizes provided insufficient power to detect subtle abnormalities.

**Objective:** To investigate microstructural white matter alterations and their relation to clinical features in the largest dataset of adult and pediatric OCD to date.

**Design, Setting, and Participants:** In this cross-sectional case-control magnetic resonance study, we investigated diffusion tensor imaging metrics from 700 adult patients and 645 adult controls, as well as 174 pediatric patients and 144 pediatric controls across 19 sites participating in the ENIGMA-OCD Working Group.

**Main Outcomes and Measures:** We extracted measures of fractional anisotropy (FA) as main outcome, and mean diffusivity, radial diffusivity, and axial diffusivity as secondary outcomes for 25 white matter regions. We meta-analyzed patient-control group differences (Cohen’s *d*) across sites, after adjusting for age and sex, and investigated associations with clinical characteristics.

**Results:** Adult OCD patients showed significant FA reduction in the sagittal stratum (*d*=-0.21, z=-3.21, p=0.001) and posterior thalamic radiation (*d*=-0.26, *z*=-4.57, p<0.0001). In the sagittal stratum only, lower FA was associated with a younger age of onset (z=2.71, p=0.006), longer duration of illness (z=-2.086, p=0.036) and a higher percentage of medicated patients in the cohorts studied (z=-1.98, p=0.047). No significant association with symptom severity was found. Pediatric OCD patients did not show any detectable microstructural abnormalities compared to matched controls.

**Conclusions and Relevance:** Microstructural alterations in projection and association fibers to posterior brain regions were found in adult OCD, and related to disease course and medication status. Such results are relevant to models positing deficits in connectivity as a crucial mechanism in OCD.

**KEY POINTS:** *Question:* Do patients with Obsessive-Compulsive Disorder (OCD) show white matter microstructural alterations, and are these alterations related to clinical features?

*Findings:* Data from 19 sites of the ENIGMA-OCD Consortium were included, involving 700 adult patients and 645 adult controls, 174 pediatric patients and 144 pediatric controls. Diffusion tensor imaging data were meta-analyzed using a harmonized data processing and analysis protocol. Adult, but not pediatric, patients showed alterations in the sagittal stratum and posterior thalamic radiation; sagittal stratum differences were associated with clinical features.

*Meaning:* Microstructural abnormalities found in adult but not in the pediatric cohort, are related to illness duration and medication status.

## INTRODUCTION

Abnormalities in cerebral white matter (WM) are relevant to models of anomalous brain circuitry that posit deficits in connectivity in obsessive– compulsive disorder (OCD)^1,2^. OCD has a childhood onset in over 50% of all cases, and most childhood-onset OCD cases persist into adulthood^3^. Diffusion tensor imaging (DTI) allows the study of WM at the microstructural level through the analysis of intrinsic, three-dimensional diffusion properties of water within brain tissues^4^. Prior DTI studies in OCD^5–7^ suggest that microstructural alterations are present in a number of WM areas. However, results across studies are inconsistent, with contrasting or conflicting effects of OCD on DTI metrics^8^. Sources of heterogeneity may include methodological factors (e.g., imaging acquisition and data processing), clinical characteristics, and variations in demographic or socioeconomic factors. More importantly, sample size variations may impact reported findings, as small studies may have insufficient power to detect subtle alterations^9^. Brain imaging consortia offer new opportunities, pooling data and findings from around the world to achieve an appropriate sample size. The OCD working group of the Enhancing Neuro-Imaging Genetics through Meta-Analysis (ENIGMA) consortium^10^, is one such collaboration. Previous findings from the working group focused on subcortical and cortical brain grey matter abnormalities, using subcortical volumes, cortical thickness and surface area quantification algorithms. An initial analysis of data from 3,589 individuals showed distinct subcortical volume abnormalities in adults (smaller hippocampal and larger pallidal volumes) and unmedicated children (larger thalamic volume) with OCD^11^. The second study focused on cortical grey matter differences and showed a lower surface area for the transverse temporal cortex and a thinner inferior parietal cortex in adult patients. In pediatric OCD patients compared to healthy controls, significantly thinner inferior and superior parietal cortices were found^12^. Medication status was associated with structural differences in both pediatric and adult OCD.

Here we aimed to investigate WM microstructural alterations in adult and pediatric OCD using data from the ENIGMA OCD working group, in the subset of participants that had collected diffusion MRI. DTI metrics in 700 adult patients were compared to those of 645 adult controls, and separately, 174 pediatric patients were compared to 144 pediatric controls. Analyses also aimed to investigate associations between WM microstructure and demographic and clinical variables. As prior meta-analytic findings in frontal and callosal regions have been inconsistent (with either higher^1^ or lower^1,5^ fractional anisotropy -FA- in anterior midline tracts), with more homogenous findings for fronto-temporal and fronto-parietal intra-hemispheric bundles, we expected to find microstructural alterations (as reflected by lower FA^1,5,6,8^) in the long tracts connecting frontal regions to posterior temporal, parietal and occipital association cortices.

## METHODS

### Study dataset

The ENIGMA-OCD Working Group includes 19 international research institutes. Previous literature (including studies from the present Working Group^11,12^) showed different patterns of effects in pediatric and adult cohorts; thus, we performed separate meta-analyses for adult and pediatric data. Globally, we analyzed data from 1,345 adults (including 700 OCD patients and 645 controls, aged ≥18) and 318 children (including 174 OCD and 144 controls). Patients were administered the Yale-Brown Obsessive-Compulsive Scale (Y-BOCS)^13^ and the Child Yale-Brown Obsessive-Compulsive Scale (CY-BOCS)^14^ to assess symptom severity. These scales are clinician-rated, 10-item scales, with each item rated from 0 (no symptoms) to 4 (extreme symptoms; total range, 0 to 40), with separate subtotals for severity of obsessions and compulsions. **Tables 1** and **2** show the demographic and clinical characteristics of the participants from each site.

All local IRBs approved the use of measures extracted from completely anonymized data.

### Image Acquisition and Processing

Harmonized preprocessing, including brain extraction, eddy current correction, movement correction, echo-planar imaging-induced distortion correction and tensor fitting, was carried out at each site, using protocols and quality control pipelines provided by the ENIGMA-DTI working group (http://enigma.ini.usc.edu/protocols/dti-protocols/) and already employed to pool harmonized DTI analyses from around the world^15–17^.

Once tensors were estimated, each site conducted a harmonized image analysis for FA quantification using the ENIGMA-DTI protocol, consisting of the tract-based spatial statistics (TBSS)^18^ analytic method modified to project individual FA values to the ENIGMA-DTI skeleton. Tract-wise regions of interest (ROIs), derived from the Johns Hopkins University (JHU)^19^ white matter parcellation atlas, were used to extract the mean FA across the full skeleton and mean FA values for 25 ROIs.

Diffusivity measures (i.e., mean diffusivity (MD), axial diffusivity (AD) and radial diffusivity (RD)) were also derived for secondary analysis. In the main analyses, we combined left and right regions of interest (ROI) across hemispheres, as we had no *a priori* hypotheses regarding lateralized effects on FA.

### Statistical Analysis

At each site, Cohen’s *d* effect sizes were calculated for differences in FA between patients and healthy controls. Age, sex, age-by-sex interaction and quadratic covariates of age^2^ and age^2^-by-sex interaction were included in the model, as linear and nonlinear age and sex interactions have been reported for FA^15^. Subsequently, a random effects meta-analysis was run at the coordinating site using Comprehensive Meta-Analysis (CMA, version 2, Biostat, Englewood, NJ) to combine individual site effect sizes. Heterogeneity scores (I^2^; lower values indicate lower variance in the effect size estimates across studies) were also computed for each test.

Effect sizes are reported as overall Cohen’s *d* values for case/control effects and *Z*-scores for quantitative effects from linear regressions. To control the probability of false positives due to multiple comparisons, effects of FA differences between cases and controls were considered significant if they survived the Bonferroni correction threshold of 0.05/25 (the 25 considered tracts)= 0.002. The stability of the overall effect size estimate was tested using a ‘leave one out’ sensitivity analysis. This analysis shows how the overall effect size changes if one dataset at a time is removed, assessing whether potential results are site-dependent. Furthermore, to ascertain whether the estimated effect size varied as a function of clinical characteristics, mixed effects meta-regressions were performed on diffusion parameters using age, age of onset, duration of illness, illness severity and percentage of medicated patients in the patients’ dataset as regressors. The influence of medication status was also explored through a mixed-effects sub-group analysis, comparing effect sizes in medicated (n=8) and unmedicated (n=3) patient cohorts.

## RESULTS

Demographics and clinical characteristics of the participants in each site are shown in Tables 1 and 2.

### Adult Cohort

Table 3 indicates the five out of 25 regions with marginally lower FA in patients compared to controls. These are the *genu* of the corpus callosum, the posterior *corona radiata*, the posterior thalamic radiation (PTR), the sagittal stratum (SS) and the uncinate fasciculus (see Figure 1).

**Figure 1:**
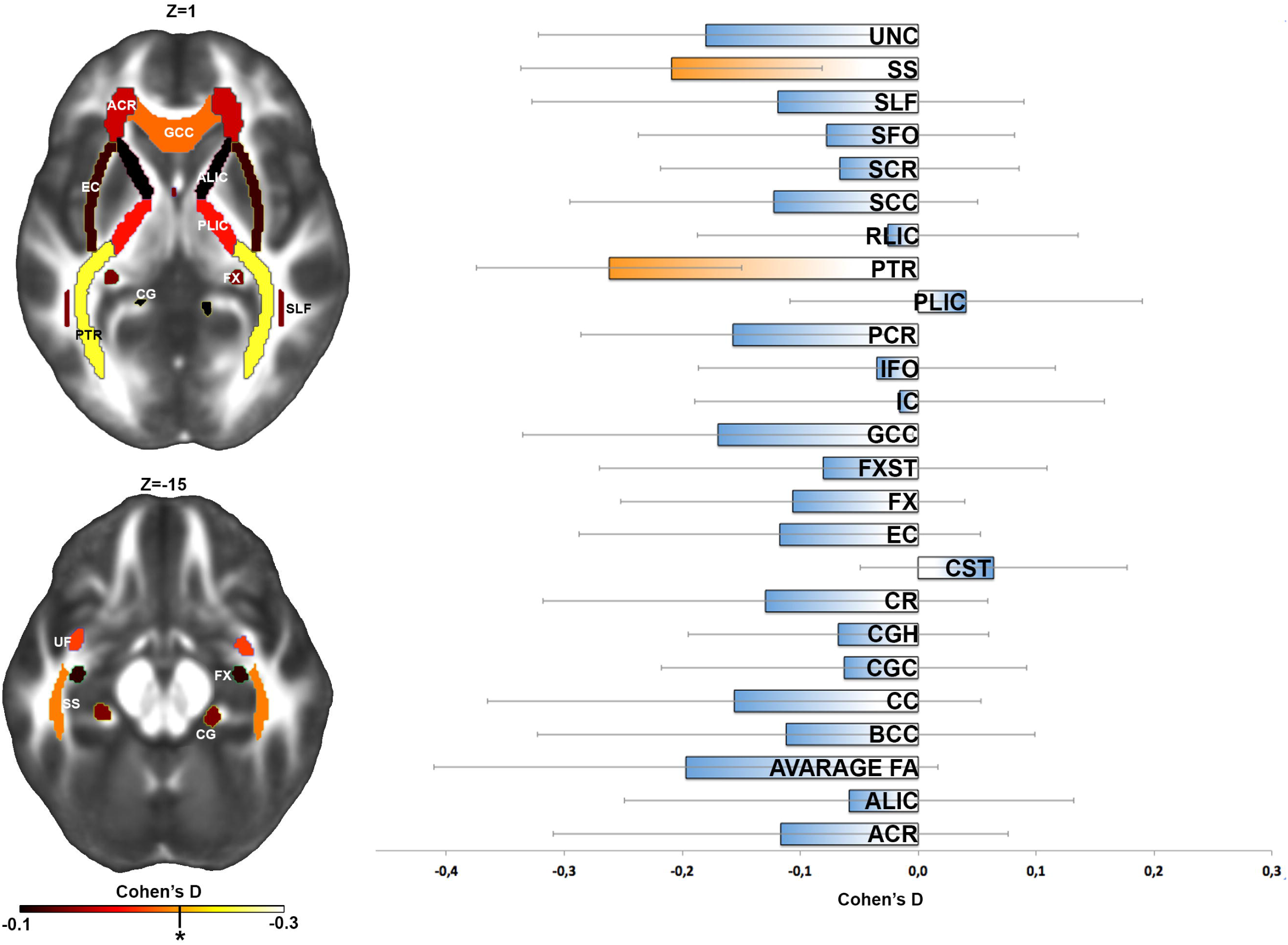
Left Panel - Fractional anisotropy (FA) differences between OCD patients and healthy controls for 25 white matter (WM) regions. Gradient bar indicates Cohen’s *d* effect sizes after meta-analysis. * Threshold for significance after correction for multiple comparisons using Bonferroni correction (P.05/25 =0.002). Right Panel-Cohen’s *d* effect sizes after meta-analysis, after including age, sex, age × sex, age^2^ and age^2^ × sex as covariates. Error bars represent 95% confidence intervals. Significant regions after adjusting for multiple regions tested are highlighted in orange.

FA in the SS (d=-0.21, z=-3.21, p=0.001) and the PTR (d=-0.261, z=-4.57, p<0.0001) survived correction for multiple comparisons. For these two regions, heterogeneity values were the lowest and did not reach statistical significance (I^2^=24.43, p=0.21 and I^2^=6.61, p=0.38, respectively) implying that effect size estimates across sites were highly consistent for regions where results survived correction. A sensitivity analysis for the PTR showed that the OCD effect was robust, yet, for the SS, Bonferroni-corrected significance was lost after removing single datasets. However, when more homogenous cohorts (i.e., exclusively medicated patients) were considered, the effect of OCD diagnosis remained significant across combinations of sites.

We also analyzed diffusivity measures (i.e., MD, AD and RD) in the entire dataset. OCD patients showed higher RD in the PTR and SS, but neither result survived the Bonferroni correction threshold (d=0.16, p=0.002 for PTR and d=0.21, p=0.007 for SS). No significant results were found for MD or AD. Meta-regressions were run in regions where a significant effect of group emerged (i.e., PTR and SS). In the SS of adults diagnosed with OCD, lower FA was significantly associated with younger age of onset (z=2.71, p=0.006), longer duration of illness (z=-2.09, p=0.036) and a higher percentage of medicated patients (z=-1.98, p=0.047; see Figure 2). Mixed effects sub-groups analysis showed a significant difference (Q-value_(df=1)_=5.27, p=0.022) between the effect sizes in medicated (N=544, d=-0.274, p=<.0001) and unmedicated (N=158, d=0.046, p=0.72) patients. No relationship was found between FA values of the PTR and clinical measures. No relationship was found between FA and YBOCS scores.

**Figure 2:**
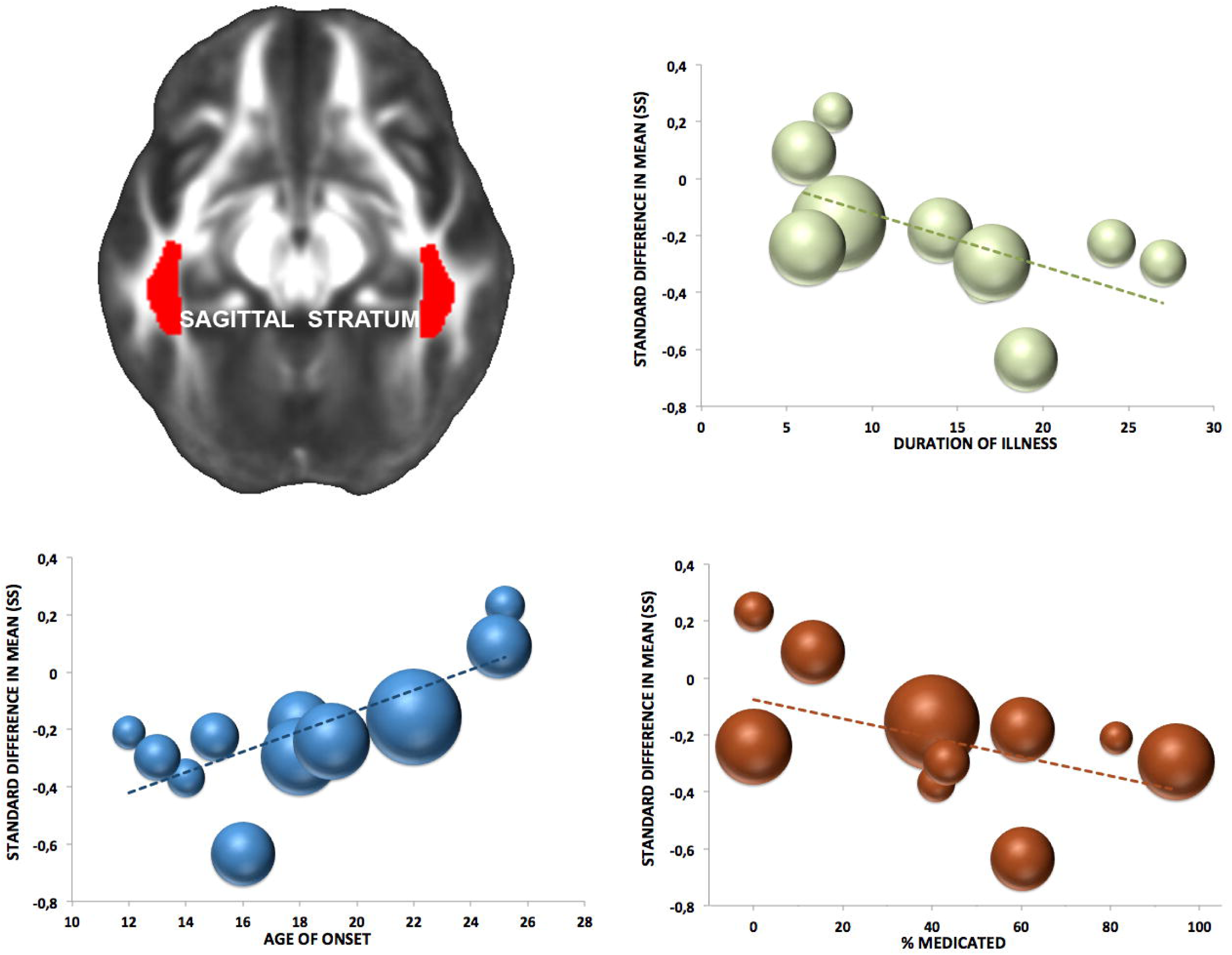
Association between FA reduction (in OCD adult patients) in the sagittal stratum and illness duration (*green dots*), age of onset (*blue dots*) and percentage of medicated patients across the 11 ENIGMA-OCD sites. Sphere magnitude indicates sample size.

### Pediatric Cohort

In the pediatric cohort, patients showed no detectable FA abnormalities in any of the regions studied (see Table 4 for statistical details).

## DISCUSSION

In the largest coordinated meta-analysis of WM in OCD to date, we demonstrated specific regional WM alterations, in adults with OCD with lower FA in the posterior thalamic radiation (PTR) and sagittal stratum (SS). Meta-regression indicated that lower FA in the SS is associated with younger age at onset, longer duration of illness, and being on medication, but not with symptom severity - implying that, as with cortical thickness and subcortical volumes in OCD, the observed alterations may be markers of the disorder. Findings were significant across the combinations of datasets in the sensitivity analysis for the PTR, while for the SS the effect remained significant when solely medicated patients were considered. We did not find case-control differences in WM microstructure of pediatric subjects.

A role for cerebral WM and oligodendrocytes (the myelinating cells of the central nervous system) in the pathophysiology of many psychiatric disorders has been supported by growing research evidence^6,20^, suggesting abnormalities of myelination status as a possible pathogenic mechanism^21^. Specifically, altered myelin-related maturational growth may explain the enhanced risk for psychiatric disorders during the transition from childhood to adulthood^22,23^, an age window of intense ongoing brain development^21^.

Although FA is a general measure of microstructure - including variation in regional myelination levels, such as axon demyelination or loss, myelin loss or increased extracellular space - it does not provide a physiologically specific explanation of WM abnormalities^24^. In our study, higher RD in the same bundles, although at a trend level, supports the hypothesis that lower FA reflects a disruption of myelin sheaths^25^, given that RD is a putative myelin marker^24^. The association between myelin degradation in the SS and longer illness duration together with the absence of a detectable alteration in pediatric patients, suggests that neuroplastic changes are epiphenomena of prolonged symptomatology, since compulsively engaging in a particular behavior or cognitive process has been suggested to alter brain structure^26,27^. Moreover, symptoms indicative of obsessive–compulsive traits are related to individual myelination over time even in otherwise healthy samples. This suggests that the mechanisms underlying compulsivity have long-lasting effects on brain development, possibly affecting myelination trajectories during adolescence, with lasting effects into adulthood^21^. Alternatively, the paucity of extrinsic factors regulating the development of myelinating glia^28^ could have driven the altered myelination and its association with prolonged illness. Indeed, impoverished environment is both the cause and the consequence of mental illness in general, and of compulsivity in particular^29^.

Drug-induced reductions in the FA of several WM tracts may be part of the mechanism of action of drug treatments in OCD^30^. Long-term treatment exposure may negatively influence the proliferation of oligodendrocytes and their myelination of axons^25,31–33^. Drug treatments for psychiatric disorders may act in part through effects on myelinating glia, as oligodendrocytes have neurotransmitter receptors for glutamate, serotonin, and dopamine. Drugs acting through these neurotransmitter systems can exert actions on myelination that may be detrimental^30,33^ or beneficial^34^. In the present study, medication effects could not be explored in great detail, but lower FA in the SS was related to medication status and observed only in the medicated patient cohort. Moreover, the effect remained significant across combinations of datasets when only medicated patients were considered, strengthening the hypothesis that medication impacts WM microstructure.

Although the mechanisms of altered WM microstructure in OCD are far from clear, a number of lines of research^35,36^ and prior genetic association studies have described specific genes related to myelination^37^, while there is evidence that genetic variants related to glutamatergic, dopaminergic, and neurodevelopmental pathways determine WM microstructure in children and adolescents with OCD^38^. Although no genes related to myelination have yet reached genome wide-significance, given preliminary findings from small samples, it is possible that such significance will be reached with better powered GWASs. Particularly, variants in the transcription factor of the oligodendrocyte lineage (OLIG2), involved in myelination and neurogenesis and essential in regulating the development of myelin producing cells, and in the myelin oligodendrocyte glycoprotein (MOG), implicated in maintaining myelin-axon integrity in the normal brain, have been associated with OCD in familial and association studies^39–41^. Although not at a genome-wide level of evidence, a single nucleotide polymorphism in the OLIG2 gene (related to psychiatric disorders including OCD^40^) was associated with reduced FA in WM regions of healthy subjects^42^, suggesting that it may confer increased risk for psychiatric illnesses through an effect on neuroanatomical connectivity. The additional observation that polymorphisms of genes codifying for myelin-promoting factors like the brain-derived neurotrophic factor (BDNF) are associated with OCD (specifically with early onset)^43^ and particularly with WM microstructure^38^, further suggests a role for altered myelination in the pathophysiology of the disorder. However, such interpretative framework of genetic influence on altered myelination in OCD is rather speculative and should be considered with caution, given that the cited studies are underpowered.

Axon-derived signals, like glutamate release, may also play a role in the epigenetic regulation of the transcriptional apparatus required for myelination by oligodendrocytes^44^, and in the process of remyelination after damage. Glutamate-related genes are promising candidates for OCD^45–47^, and neuroimaging-gene association studies in animal models reinforce the hypothesis of glutamate involvement in the pathophysiology of the disorder^48^. Since abnormalities in glutamate neurotransmission and homeostasis have shown to be crucial for OCD onset^49^, glutamate excitotoxicity-induced damage of the myelin sheaths may further explain the decline in myelin integrity suggested here. The PTR and, to a lesser extent, the SS convey fibers principally from the thalamus, a region found to be consistently enlarged in unmedicated pediatric OCD samples^11^ and functionally hyperactive^50^ in OCD adults. This pathological hyperactivity could increase glutamate release^51^, and excess glutamate from thalamic projections may determine the microstructural alteration in WM bundles reported here. Several lines of evidence from magnetic resonance spectroscopy (MRS) and from clinical trials support the existence of glutamatergic hyperactivity associated with OCD^51^. This indicates that the disruption of microstructure in the PTR and SS may partially be a consequence of other processes (e.g., glutamate abnormalities, a potential epiphenomenon of neuronal hyperactivity in OCD circuitry) in the pathophysiology of OCD^51^.

Since both the PTR and the SS convey projection fibers to the posterior part of the brain, our results support the idea that OCD involves abnormalities affecting a network of regions that is more extensive than commonly believed^1,52^. Both bundles project to posterior parietal, temporal and occipital cortices, and include many major association fibers (i.e., the inferior longitudinal fasciculus and the inferior fronto-occipital fasciculus). Their altered microstructure may be related to the cognitive dysfunctions subtended by intra-hemispheric disconnection^2,53^.

Notably, in our study WM microstructural alterations in OCD were associated with age at scanning. Specifically, WM alterations were observed in the adult cohort only, and were associated with longer illness duration. These findings, which were unrelated to OCD symptom severity, complement previous evidence of differences between adult and pediatric OCD patients in brain morphological^5,12,54^ and clinical^55^ correlates.

There is evidence that the human brain’s protracted myelination^56,57^ underpins myelin vulnerability along a continuum from early to late stages of development and disease^58^. Thus, it has been suggested that pediatric OCD could be a neurodevelopmental disorder with potentially differing patterns of myelination occurring throughout life^59^. Indeed, evidence in healthy subjects indicates that the psychiatric trait of compulsivity is linked to reduced myelin growth that emerges only during adolescence (being present only to a minor extent in childhood) as a result of aberrant developmental processes^21^. Intriguingly, MOG influence on myelin maturation and remodeling is different depending on the age of onset^60^ in those psychiatric disorders with microstructural white matter abnormalities overlapping with OCD^6^. Thus, brain changes associated with OCD genetic vulnerability may be dynamic over time, and dependent on timing of gene expression^61^, neurodevelopmental stage at illness onset, and disease course afterwards. Alternatively, pediatric OCD might be a developmentally moderated expression of etiologic processes that are shared with the adult clinical phenotype.

A number of limitations of the data analyzed here deserve emphasis. First, although TBSS is a widely used method for voxel-based analysis of WM, addressing issues associated with smoothing and misalignment in DTI group analysis^52^, the technique has some limitations. Indeed, by reducing WM tracts into a skeleton, delineating the center of the tracts and projecting onto it only the highest FA value along the projection, some information might be lost^62^ and potential artifacts, resulting from misregistration, might be produced^63^. Nevertheless, several test–retest and reliability analyses were conducted by the ENIGMA-DTI working group to ensure reproducibility of measures and effects using this TBSS approach^64^. Moreover, a word of caution is needed regarding the interpretation of the neurobiological basis of DTI measures since although FA reflects the myelination, orientational coherence, and microtubular axonal structure of fibers, other in vivo markers not explored in the present study have been shown to be a more direct reflection of myelination status^65,66^. Another potential limitation of the present study may lie in the differences in clinical characteristics between the studied patients, particularly in the average age of onset (which ranged from 4 to 49). Since the latter is often calculated retrospectively, a reliable and unanimous method for establishing this important effect moderator is warranted. Also, we were not able to calculate specific dosages of different medication types and analyze medication effects in terms of drug dosages or total time of treatment. Thus, our interpretation that a disruption of myelin sheaths in the PTR and SS may be a downstream consequence of prolonged medication or glutamatergic abnormalities in OCD should be viewed with caution, as the potential detrimental/normalizing effects of different medications could not be fully tested. Lastly, it is worth mentioning that while the adult cohort analysis had sufficient power to detect the observed effect size, as the sample size was adequate to detect microstructural differences as small as d=0.15, the null result in the pediatric cohort may be a consequence of the relatively small sample size since the power for potentially detecting even very small differences was low (0.32). Nevertheless, this is the largest pediatric dataset investigated in a DTI study of OCD.

In summary, our results clearly indicate a key role in adult OCD of microstructural alterations in projection and association fibers to posterior brain regions. They also demonstrate the moderating effect of illness chronicity and medication on WM microstructure. Such alterations might be a downstream consequence of other processes that are more primary to the pathophysiology of the disorder and potentially related to OCD genetic vulnerability. Our meta-regression results related to duration suggest that microstructural alterations may persist along the course of the illness, although longitudinal data will be needed to confirm such trajectories. Thus, from a clinical perspective, early symptom reduction and subsequent improved functioning might counteract genetic vulnerability and promote the epigenetic regulation of myelin remodeling^67,68^ by enhancing extrinsic factors mediating myelin plasticity^28^. However, pharmacological amelioration of symptoms should be cautiously considered, given the detrimental medication effect we observed on WM microstructure. Other therapeutic approaches, such as cognitive-behavioural^69^ therapies should be strongly encouraged.

Future studies to investigate the co-occurrence of abnormal WM microstructure, GM volume and metabolic differences in OCD will shed light on the interactions and trajectories of structural and functional alterations in this condition. In particular, longitudinal designs, and collecting information from patients at their illness onset, combined with multimodal MRI approaches such as volumetric, DTI, fMRI, and MRS will help provide an understanding of the timing and course of brain changes in OCD and provide greater insight into the mechanisms involved in various stages of OCD, including the long-term effects of medication.

