## Supplementary material for "White Matter Microstructure and its Relation to Clinical Features of Obsessive-Compulsive Disorder: Findings from the ENIGMA OCD Working Group": Table 1

| **Charasteristic** | **Adult OCD Sample**  **(n=700)** | **Adult HC Sample**  **(n=645)** | **Pediatric OCD Sample**  **(n=174)** | **Pediatric HC Sample**  **(n=144)** |
| --- | --- | --- | --- | --- |
| Age (years) | 31.4 ± 9.9 | 30.7±10 | 14.5 ± 2.3 | 14.3 ± 2.5 |
| OCD illness severity score | 25 ± 7.1 | - | 20.7 ± 7.8 | - |
| Age at onset | 19.1 ± 8.4 | - | 13.1 ± 5.3 | - |
|  | N (%) | N (%) | N (%) | N (%) |
| Male | 405 (58) | 378 (59) | 94 (53) | 74 (51.3) |
| Medication use at time of scan | 269 (39) | - | 112 (64) | - |
| Current comorbid disorders |  |  |  |  |
| Anxiety | 74^a^ (11) | - | 27^d^ (23) | - |
| Major depression | 60^b^ (10) | - | 10^d^ (9) | - |
| OCD symptom dimension |  |  |  |  |
| Aggressive/Checking | 418^c^(79) | - | 82^e^ (75) | - |
| Contamination/cleaning | 358^c^(68) | - | 74^e^ (68) | - |
| Symmetry/ordering | 378^c^(62) | - | 72^e^ (66) | - |
| Sexual/religious | 229^c^(44) | - | 47^e^ (43) |  |
| Hoarding | 115^c^(22) | - | 45^e^ (41) | - |

TABLE 1: Demographic and Clinical Characteristics of Patients With Obsessive-Compulsive Disorder (OCD) and Control Subjects

^a^Data available for 625 patients; ^b^data available for 610 patients; ^c^data available for 525 patients; ^d^data available for 117 patients; ^e^data available for 109 patients
