## Supplementary material for "White Matter Microstructure and its Relation to Clinical Features of Obsessive-Compulsive Disorder: Findings from the ENIGMA OCD Working Group": Table 2

Table 2: Breakdown, by Site, of Clinical Characteristics of Patients With Obsessive-Compulsive Disorder (OCD) in the ENIGMA OCD

Working Group Samples

| **Site** | **Medicated (%)** | **Age of Onset** | **Duration of illness** | **YBOCS score** | **Lifetime Anxiety (%)** | **Lifetime Depression %** |
| --- | --- | --- | --- | --- | --- | --- |
| Amsterdam | 0 | 15,1±6,8 | 23,7±12,8 | 21,3±6,1 | 42,1 | 47,4 |
| Bangelore | 39,9 | 22,3±7,7 | 7,2±5,2 | 25,5±6,5 | 8,9 | 7 |
| Capetown | 40,9 | 13,2±5,7 | 17,2±11,5 | 23±4,2 | 0 | 0 |
| Kyoto | 0 | 25,2±9 | 7,7±6,2 | 21,9±6,6 | 8,6 | 0 |
| Milan | 60,3 | 15,6±6,2 | 18,9±11,6 | 31,4±5,2 | 1,6 | 7,9 |
| Mount Sinai | 81,3 | 12,4±5,7 | 15,1±6,7 | 19,9±5,9 | 50 | 18,8 |
| Munich | 60,3 | 17,6±6,7 | 13,5±10,3 | 20,8±6,2 | 8,2 | 23,3 |
| Rome | 94,8 | 16,8±8 | 17,3±12,8 | 23,2±9,3 | 10,4 | 9,1 |
| Sao Paulo | 43,2 | 12,8±5,9 | 26,3±13,5 | 29,2±6,2 | 73 | 83,8 |
| Shangai | 0 | 24±9,9 | 6±5,9 | 26,2±4,7 | 0 | 0 |
| Seoul | 13,3 | 19,1±7,2 | 6,2±7 | 25,8±6,9 | 1 | 2 |

Legenda: YBOCS=Yale-Brown Obsessive Compulsive Scale
