## Supplementary material for "White Matter Microstructure and its Relation to Clinical Features of Obsessive-Compulsive Disorder: Findings from the ENIGMA OCD Working Group": Table 3

|  |  | **Effect size and 95% confidence interval** | | | | | | | | |  | | **Heterogeneity** | | | | | |
| --- | --- | --- | --- | --- | --- | --- | --- | --- | --- | --- | --- | --- | --- | --- | --- | --- | --- | --- |
| **ROI** |  | **Cohen's d** | **S.E.** | **Lower limit** | | **Upper limit** | | **Z-value** | | **P-value** | |  | | **Q-value** | | **P-value** | | **I-squared** |
| ACR |  | -0,1164 | 0,0983 | -0,3091 | 0,0763 | | -1,1839 | | 0,2364 | |  | | 29,1988 | | 0,0012 | | 65,7520 | |
| ALIC |  | -0,0584 | 0,0971 | -0,2488 | 0,1320 | | -0,6013 | | 0,5477 | |  | | 28,5200 | | 0,0015 | | 64,9369 | |
| AverageFA |  | -0,1968 | 0,1091 | -0,4107 | 0,0171 | | -1,8036 | | 0,0713 | |  | | 35,9099 | | 0,0001 | | 72,1525 | |
| BCC |  | -0,1119 | 0,1076 | -0,3227 | 0,0990 | | -1,0398 | | 0,2984 | |  | | 34,9932 | | 0,0001 | | 71,4230 | |
| CC |  | -0,1558 | 0,1067 | -0,3650 | 0,0533 | | -1,4606 | | 0,1441 | |  | | 34,4018 | | 0,0002 | | 70,9317 | |
| CGC |  | -0,0626 | 0,0789 | -0,2173 | 0,0920 | | -0,7938 | | 0,4273 | |  | | 18,9453 | | 0,0410 | | 47,2164 | |
| CGH |  | -0,0677 | 0,0650 | -0,1951 | 0,0598 | | -1,0404 | | 0,2982 | |  | | 13,2641 | | 0,2093 | | 24,6083 | |
| CR |  | -0,1294 | 0,0962 | -0,3179 | 0,0591 | | -1,3454 | | 0,1785 | |  | | 27,9259 | | 0,0019 | | 64,1910 | |
| CST |  | 0,0641 | 0,0577 | -0,0490 | 0,1772 | | 1,1106 | | 0,2667 | |  | | 10,8843 | | 0,3666 | | 8,1241 | |
| EC |  | -0,1173 | 0,0868 | -0,2873 | 0,0528 | | -1,3513 | | 0,1766 | |  | | 22,7693 | | 0,0116 | | 56,0812 | |
| FX |  | -0,1063 | 0,0745 | -0,2523 | 0,0398 | | -1,4259 | | 0,1539 | |  | | 16,9566 | | 0,0753 | | 41,0259 | |
| FXST |  | -0,0804 | 0,0968 | -0,2701 | 0,1093 | | -0,8307 | | 0,4062 | |  | | 28,3405 | | 0,0016 | | 64,7148 | |
| GCC |  | -0,1696 | 0,0845 | -0,3352 | -0,0041 | | -2,0085 | | 0,0446 | |  | | 21,5584 | | 0,0175 | | 53,6144 | |
| IC |  | -0,0158 | 0,0887 | -0,1896 | 0,1581 | | -0,1776 | | 0,8590 | |  | | 23,8155 | | 0,0081 | | 58,0106 | |
| IFO |  | -0,0350 | 0,0772 | -0,1864 | 0,1164 | | -0,4531 | | 0,6505 | |  | | 18,1854 | | 0,0519 | | 45,0108 | |
| PCR |  | -0,1570 | 0,0660 | -0,2863 | -0,0277 | | -2,3803 | | 0,0173 | |  | | 13,5590 | | 0,1941 | | 26,2484 | |
| PLIC |  | 0,0406 | 0,0763 | -0,1090 | 0,1903 | | 0,5321 | | 0,5947 | |  | | 17,7645 | | 0,0591 | | 43,7078 | |
| PTR |  | -0,2619 | 0,0573 | -0,3742 | -0,1495 | | -4,5689 | | 0,0000 | |  | | 10,7084 | | 0,3807 | | 6,6151 | |
| RLIC |  | -0,0256 | 0,0823 | -0,1869 | 0,1356 | | -0,3117 | | 0,7553 | |  | | 20,5308 | | 0,0246 | | 51,2926 | |
| SCC |  | -0,1223 | 0,0882 | -0,2952 | 0,0506 | | -1,3868 | | 0,1655 | |  | | 23,5299 | | 0,0090 | | 57,5008 | |
| SCR |  | -0,0664 | 0,0776 | -0,2184 | 0,0856 | | -0,8565 | | 0,3917 | |  | | 18,3052 | | 0,0500 | | 45,3707 | |
| SFO |  | -0,0776 | 0,0813 | -0,2370 | 0,0818 | | -0,9545 | | 0,3399 | |  | | 20,0600 | | 0,0287 | | 50,1495 | |
| SLF |  | -0,1189 | 0,1066 | -0,3278 | 0,0901 | | -1,1149 | | 0,2649 | |  | | 34,3842 | | 0,0002 | | 70,9169 | |
| SS |  | -0,2090 | 0,0651 | -0,3367 | -0,0814 | | -3,2100 | | 0,0013 | |  | | 13,2328 | | 0,2109 | | 24,4304 | |
| UNC |  | -0,1799 | 0,0723 | -0,3216 | -0,0382 | | -2,4885 | | 0,0128 | |  | | 15,9772 | | 0,1003 | | 37,4109 | |

TABLE 3: FA meta-analysis metrics for the adult sample. Cohen’s d values, their s.e., Lower and Upper limits, P-values and I^2^ (heterogeneity) values after meta-analysis for differences between OCD patients and healthy controls
