## Supplementary material for "White Matter Microstructure and its Relation to Clinical Features of Obsessive-Compulsive Disorder: Findings from the ENIGMA OCD Working Group": Table 4

|  |  | **Effect size and 95% confidence interval** | | | | | | | | |  | | **Heterogeneity** | | | | | |
| --- | --- | --- | --- | --- | --- | --- | --- | --- | --- | --- | --- | --- | --- | --- | --- | --- | --- | --- |
| **ROI** |  | **Cohen's d** | **S.E.** | **Lower limit** | | **Upper limit** | | **Z-value** | | **P-value** | |  | | **Q-value** | | **P-value** | | **I-squared** |
| ACR |  | 0,035 | 0,111 | -0,183 | 0,254 | | 0,318 | | 0,751 | |  | | 4,154 | | 0,762 | | 0,000 | |
| ALIC |  | 0,103 | 0,145 | -0,180 | 0,386 | | 0,711 | | 0,477 | |  | | 11,207 | | 0,130 | | 37,538 | |
| AverageFA |  | 0,077 | 0,111 | -0,142 | 0,295 | | 0,687 | | 0,492 | |  | | 5,421 | | 0,609 | | 0,000 | |
| BCC |  | 0,011 | 0,138 | -0,260 | 0,282 | | 0,079 | | 0,937 | |  | | 10,316 | | 0,171 | | 32,143 | |
| CC |  | -0,013 | 0,117 | -0,242 | 0,217 | | -0,108 | | 0,914 | |  | | 7,596 | | 0,370 | | 7,844 | |
| CGC |  | 0,111 | 0,112 | -0,107 | 0,330 | | 0,997 | | 0,319 | |  | | 6,212 | | 0,515 | | 0,000 | |
| CGH |  | -0,017 | 0,143 | -0,296 | 0,263 | | -0,116 | | 0,908 | |  | | 10,928 | | 0,142 | | 35,946 | |
| CR |  | 0,022 | 0,111 | -0,196 | 0,240 | | 0,197 | | 0,844 | |  | | 5,042 | | 0,655 | | 0,000 | |
| CST |  | -0,110 | 0,153 | -0,409 | 0,189 | | -0,723 | | 0,469 | |  | | 12,410 | | 0,088 | | 43,593 | |
| EC |  | -0,025 | 0,111 | -0,244 | 0,193 | | -0,226 | | 0,821 | |  | | 5,000 | | 0,660 | | 0,000 | |
| FX |  | -0,119 | 0,111 | -0,337 | 0,099 | | -1,072 | | 0,284 | |  | | 1,237 | | 0,990 | | 0,000 | |
| FXST |  | -0,095 | 0,134 | -0,357 | 0,168 | | -0,707 | | 0,480 | |  | | 9,725 | | 0,205 | | 28,020 | |
| GCC |  | -0,018 | 0,111 | -0,236 | 0,201 | | -0,158 | | 0,875 | |  | | 4,635 | | 0,704 | | 0,000 | |
| IC |  | 0,013 | 0,111 | -0,205 | 0,231 | | 0,116 | | 0,907 | |  | | 3,551 | | 0,830 | | 0,000 | |
| IFO |  | -0,025 | 0,111 | -0,243 | 0,193 | | -0,228 | | 0,819 | |  | | 2,937 | | 0,891 | | 0,000 | |
| PCR |  | 0,078 | 0,116 | -0,149 | 0,305 | | 0,676 | | 0,499 | |  | | 7,443 | | 0,384 | | 5,958 | |
| PLIC |  | 0,026 | 0,111 | -0,192 | 0,244 | | 0,237 | | 0,813 | |  | | 2,079 | | 0,955 | | 0,000 | |
| PTR |  | -0,004 | 0,201 | -0,398 | 0,390 | | -0,022 | | 0,982 | |  | | 21,204 | | 0,003 | | 66,987 | |
| RLIC |  | -0,067 | 0,111 | -0,285 | 0,151 | | -0,602 | | 0,547 | |  | | 5,021 | | 0,657 | | 0,000 | |
| SCC |  | -0,036 | 0,112 | -0,255 | 0,183 | | -0,321 | | 0,748 | |  | | 6,867 | | 0,443 | | 0,000 | |
| SCR |  | -0,040 | 0,111 | -0,258 | 0,178 | | -0,357 | | 0,721 | |  | | 4,165 | | 0,761 | | 0,000 | |
| SFO |  | 0,100 | 0,155 | -0,203 | 0,404 | | 0,649 | | 0,516 | |  | | 12,757 | | 0,078 | | 45,127 | |
| SLF |  | -0,089 | 0,116 | -0,317 | 0,139 | | -0,763 | | 0,446 | |  | | 7,519 | | 0,377 | | 6,904 | |
| SS |  | 0,006 | 0,146 | -0,280 | 0,292 | | 0,041 | | 0,967 | |  | | 11,433 | | 0,121 | | 38,775 | |
| UNC |  | 0,053 | 0,119 | -0,181 | 0,286 | | 0,441 | | 0,659 | |  | | 7,840 | | 0,347 | | 10,711 | |

TABLE 4: FA meta-analysis metrics for the pediatric sample. Cohen’s d values, their s.e., Lower and Upper limits, P-values and I^2^ (heterogeneity) values after meta-analysis for differences between OCD patients and healthy controls
